# Relationship between weapon size and six key behavioural and physiological traits in males of the European earwig

**DOI:** 10.1101/2024.03.20.585871

**Authors:** Samantha E.M. Blackwell, Laura Pasquier, Simon Dupont, Séverine Devers, Charlotte Lécureuil, Joël Meunier

**Affiliations:** Institut de Recherche sur la Biologie de l’Insecte (IRBI), UMR CNRS 7261, Université de Tours, France

**Keywords:** Behaviour, Insect, Metarhizium, Ornament, Sexual selection, Weapons

## Abstract

In many animals, male weapons are large and extravagant morphological structures that typically enhance fighting ability and reproductive success. It is generally assumed that growing and carrying large weapons is costly, thus only males in the best condition can afford it. In the European earwig, males carry weapons in the form of forceps-like cerci, which can vary widely in size within populations. While long forceps appear to increase male’s access to females, it is unknown whether it also correlates with other important male lifen-history traits. This information is important, however, in determining the potential reliability of forceps length as an indicator of male quality and the stability of this signalling system. Here, we tested whether forceps length is associated with six important behavioural and physiological traits in males of the European earwig. We sampled hundreds of males from two populations, selected 60 males with the longest and shortest forceps from each population, and then measured locomotor performance, boldness, aggregation behaviour, survival under harsh conditions, sperm storage, and survival after pathogen exposure. Contrary to our predictions, we detected no main association between forceps length and the traits measured. This lack of association was consistent between the two populations, although there were population-specific levels of boldness, aggregation and survival in harsh conditions (for long-forceps males only). Overall, these results challenge our current understanding of the function and quality signal of forceps length in this species and raise questions about the evolutionary drivers that could explain the maintenance of weapon size diversity within and between populations.

## Introduction

Animal reproduction often requires males to engage in physical competition and courtship to attract females in search of mates (Davies et al., 2012). From vertebrates to arthropods, these two needs have often led to the evolution of male weapons and ornaments through sexual selection (Emlen, 2008; McCullough et al., 2016; Goldberg et al., 2019). These weapons and ornaments are typically large and extravagant morphological structures that can grow on different parts of the male’s body, take a variety of forms (such as antlers, horns, spurs, fangs and tusks), and work to enhance the male’s fighting ability and/or attractiveness to females (Emlen, 2008). Textbook examples of this enhancement can be found in the white-tailed deer *Odocoileus virginianus*, where males growing the largest antlers are more likely to win fights with other males and have higher annual breeding success (Newbolt et al., 2017), and in the stalk-eyed flies *Cyrtodiopsis whitei* and *C. dahnanni*, where females prefer to mate with males exhibiting the longest eye stalks (Wilkinson et al., 1998).

However, not all males display extravagant weapons or ornaments (Emlen, 2008; McCullough et al., 2016; Goldberg et al., 2019), as the development and maintenance of these sexually selected traits often comes at a cost to males. This cost can arise from the fact that carrying heavy, bulky weapons (or ornaments) makes males less mobile and more visible to both predators and prey (Oufiero & Garland, 2007). For example, males with experimentally-enlarged wing spots have lower survival rates due to increased conspicuity to both visually orienting predators and visually orienting prey in the rubyspot damselfly *Hetaerina americana* (Grether, 1997). The cost of carrying large morphological structures can also arise because it may impose investment trade-offs with physiological functions ranging from metabolism to muscle development and spermatogenesis, which can be crucial for male fitness and survival (Emlen, 2001). For example, males with the largest hind leg weapon pay the highest resting metabolic rate and energy costs in the Hemiptera *Leptoscelis tricolor* (Somjee et al., 2018). Similarly, growing large mandibles come with low flight-muscle mass in the stag beetle *Cyclommatus metallifer* (Mills et al., 2016), and males who invest in extravagant sexual displays show a more rapid decline in spermatogenesis than males who invest less in these displays in the houbara bustard *Chlamydotis undulata* (Preston et al., 2011). As a result, only the males in the very best condition are expected to possibly afford the development and maintenance of extravagant weapons and ornaments (Otte & Stayman, 1979; Emlen & Nijhout, 2000). Determining the reliability of these morphological structures as indicators of the male quality is therefore crucial to understanding the stability of these signalling systems (Berglund et al., 1996).

Earwigs (Dermaptera) are textbook examples of insects with males displaying a sexually selected weapon. In this taxonomical group, females have relatively short, straight and hardened forceps-like cerci typically used to defend the clutch against predators (Meunier, 2024a), whereas males display elongated, curved and hardened forceps-like cerci (hereafter referred to as forceps) involved in mating courtship and fights against other males (Briceño & Eberhard, 1995; Walker & Fell, 2001; Kamimura, 2014). There are several lines of evidence to suggest that long forceps provide males with benefits in terms of mating success (Eberhard & Gutierrez, 1991; Tomkins & Brown, 2004; Kamimura, 2014). For example, males with long forceps are more likely to win fights with other males by squeezing them between their cerci, and thus gain better access to females in the toothed earwig *Vostox apicedentatus* (Moore & Wilson, 1993). In the European earwig *Forficula auricularia*, male forceps are also used in male-male contests as a weapon to deter competitors prior to mating (Styrsky & Rhein, 1999) or to interrupt mating individuals by non-copulating males (Forslund, 2000, 2003; Walker & Fell, 2001). Long-forceps males are also generally more aggressive, more readily accepted by females for mating and copulate longer compared to short-forceps males (reviewed in Kamimura, 2014). Although forceps are involved in male courtship (Walker & Fell, 2001), females do not seem to select their mate exclusively on the basis of forceps length (Radesäter & Halldórsdóttir, 1993; Forslund, 2000, 2003; Walker & Fell, 2001).

While the relationship between male forceps length and mating success has been well documented in earwigs (reviewed in Kamimura, 2014), it remains unclear whether having long or short forceps is associated with other important life-history traits in these males. However, this is important information to determine the potential reliability of forceps length as an indicator of male quality and the stability of this possible signalling system. The very few studies that have addressed this question focused on male immunity in the European earwig *F. auricularia* (Rantala et al., 2007; Körner et al., 2017). On the one hand, they show that males with long forceps have lower basal levels of certain components of the immune system, such as lysozyme activity and hemocyte concentrations. This suggests that having long forceps comes with an immune cost for males. On the other hand, they show no association between forceps length and other components of the immune response, such as encapsulation rate and phenoloxidase activity. Taken together, these findings suggest that the immune costs of having long forceps may be relatively limited and raise the question of whether other important life-history traits can be associated with forceps length.

Here, we investigated whether long and short forceps are associated with six important behavioural and physiological traits in males of the European earwig. We sampled hundreds of males in two natural populations of earwigs separated by 400 km and in each population, we selected the 30 males with the longest and the 30 males with the shortest forceps. We then measured their level of expression of three important behaviours, their survival rate in two distinct harsh conditions and their sperm quantity. The first behaviour was their locomotor performance (Cheutin et al., 2024), which reflects the ability of males to walk long distances (to forage, hide or find a mate) while carrying long and heavy (or short and light) appendages. The second behaviour was their likelihood to flee after a physical disturbance (i.e., boldness), which shows how males react when disturbed by a predator attack (Thesing et al., 2015). The third behaviour was their propensity to aggregate with conspecifics. This is an important parameter in this gregarious species, as adults typically live in groups of up to several hundred individuals and social isolation can have detrimental effects on their physiology (Kohlmeier et al., 2016; Van Meyel & Meunier, 2022). We also measured their survival rate in a harsh condition where they were isolated with no access to a food source for 31 days, and then their survival rate after exposure to the common entomopathogenic fungus *Metarhizium brunneum* (Vogelweith et al., 2017). Finally, we measured the level of sperm storage in seminal vesicles of each male, a parameter that is often important in the context of male-male competition for mating (Shuker & Simmons, 2014). If forceps length is a reliable signal of good quality, we predict that long forceps males would 1) have higher locomotor performance, 2) be bolder and thus less likely to flee after a simulated attack, 3) be more social and thus more likely to aggregate with conspecifics, 4) be more resilient to harsh environmental conditions and thus have a higher survival rate after 31 days of starvation, and/or would 5) have higher immune defences and thus survive better after a pathogen infection than short-forceps males. We also predict that 6) long-forceps males would produce more sperm and contain more sperm in their seminal vesicles than short forceps males. This last prediction would be consistent with the longer duration of copulation reported for the long-forceps males (Kamimura, 2014), even if this longer duration may also reflect other male mating strategies, such as mate guarding.

## Material and methods

### Animal rearing and selection

In July 2022, we field sampled 3000 males and females in a cultivated peach orchard near Valence, France (Lat 44.9772790, Long 4.9286990) and 400 males and females in a non-cultivated area at the edge of a forest near Cinais, France (Lat 47.1606970, Long 0.1763663). All these individuals belong to *Forficula auricularia* Linneaus, 1758, also called *Forficula auricularia* clade A (González-Miguéns et al., 2020). Immediately after field sampling, we distributed them by population of origin into 50 plastic terrariums (37 × 22 × 25 cm, balanced sex ratio) to homogenize nutrition, habitation, and access to mates for the males (Körner et al., 2017). Three months later (i.e., at the end of the reproductive season), we removed the females from all the terrariums to mimic their natural dispersal. One month later, for each population, we visually selected the 30 males with the longest forceps and the 30 males with the shortest forceps (Körner et al., 2017) and isolated them in individual Petri dishes (diameter 5 cm) for later use in behavioural and physiological measurements (A timeline of the experimental design and the detailed sample sizes can be found in Figure 1). Two days after isolation, we confirmed the robustness of our two forceps categories by measuring the mean of the left and right outer forceps of the 120 males. These measurements were done to the nearest 0.001 mm using the Leica Application Suite 4.5 software (Leica Microsystems, Wetzlar, Germany) on pictures taken under a binocular scope (Leica, MZ 12.5). These measurements confirmed that there was no overlap between the two male categories (Figure 2). All the remaining males and females that were not used in this experiment were involved in other experiments that are not presented here.

**Figure 1.**
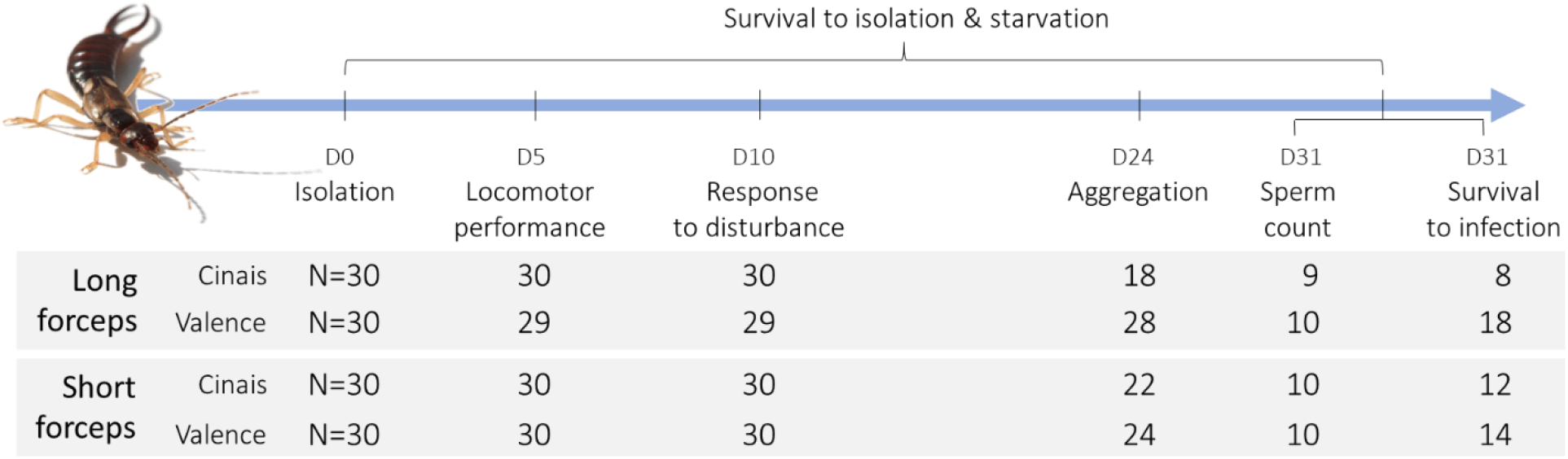
Timeline of the experimental design and evolution of sample size over the course of the experiment. The observed decrease in sample size over time reflects their mortality during isolation and in the absence of any food source. Males are distributed according to their forceps length category and population of origin.

**Figure 2.**
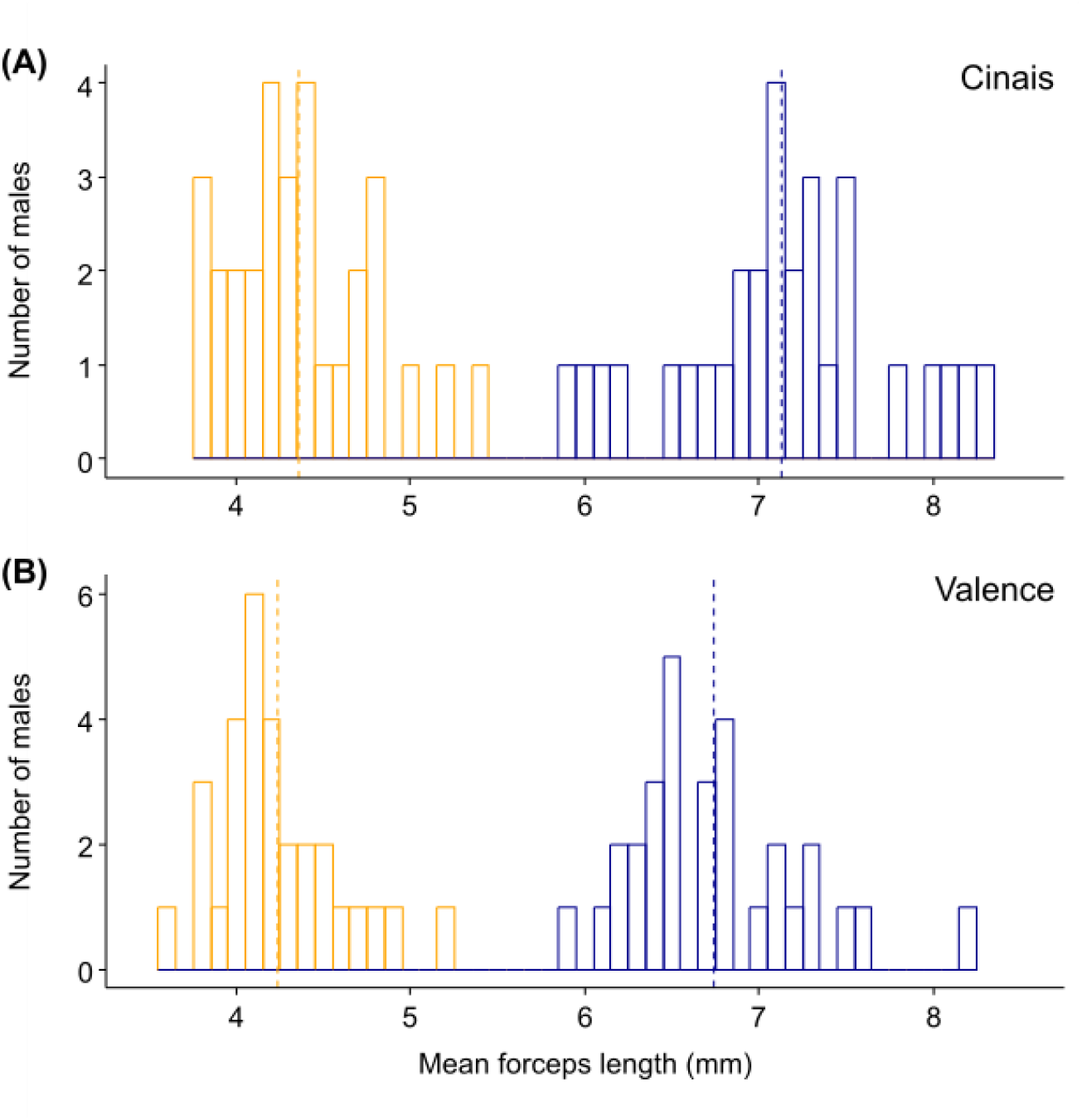
Forceps length distribution of the 120 selected males with short (orange) and long (blue) forceps in Cinais (A) and Valence (B) populations. Dashed lines show the mean values per forceps category for each population

From field sampling to isolation, we fed animals with artificial food containing mostly pollen, carrots, and cat food (see details in Kramer et al. 2015). During these three months, we offered this food *ad libitum* to homogenise the nutritional condition of the males when we started our measurements. We then kept the isolated males without access to food from the day they were isolated until 31 days after their isolation (Figure 1) to test whether resistance to both starvation and social isolation (i.e., harsh environmental conditions) were population and/or forceps-length specific, while ensuring that good rearing conditions did not mask any potential investment trade-offs between forceps length and other life history traits. We then provided food to males after day 31 to measure their survival following pathogen exposure. We kept all animals on a 12h:12h light:dark schedule at 18°C, and both the terrarium and Petri dishes contained a layer of moist sand.

### Behavioural measurements

From the 5th to the 24th day after isolation, we measured three behaviours in the 120 isolated males: the locomotor performance, the likelihood of fleeing after a physical disturbance, and the level of aggregation. All these measurements followed standard protocols for earwigs (Merleau et al., 2022; Honorio et al., 2023). First, we measured male’s locomotor performance 5 days after isolation. On that day, we transferred each male to a 3D-printed circular arena (Open field; diameter = 8 cm, height 0.4 cm) with a cover made of glass, placed on an infrared table and kept in complete darkness. We then video recorded males’ locomotion for 20 minutes (Camera: BASLER BCA 1300, Germany; Media Recorder v4.0, Noldus Information Systems, Netherland) and subsequently analyzed the resulting videos with the software Ethovision XT 16 (Noldus Information Systems, Netherland). We defined the locomotor performance of each male as the total distance he walked (in cm) during the entire recording (Merleau et al., 2022; Honorio et al., 2023).

Second, we measured the likelihood of fleeing after a physical disturbance 10 days after isolation. On that day, we carefully opened each Petri dish, pricked the male on the pronotum with a glass capillary and then recorded whether or not the male’s first reaction was to move more than one body length away from its initial position (i.e., flee).

Finally, we measured the level of aggregation of each male 24 days after isolation. We placed each male in a 3D-printed arena used in Van Meyel & Meunier, 2022, consisting of four linearly aligned circular chambers (diameter 4 cm). Three of the chambers were connected by 0.5 cm wide corridors allowing earwigs to move between chambers. The width of the corridor connecting to the fourth (outer) chamber was reduced to 0.15 cm, which prevented earwig movement while allowing the circulation of odours and antennal contacts between individuals on both sides (Van Meyel & Meunier, 2022). We started the experiment by placing two naive males and one female from the same population (but not involved in any other experiment or measurement) in this isolation chamber, and the tested male in one of the connected chambers. We recorded whether the tested male was in the chamber next to the group of conspecifics (yes or no) and repeated this measurement by taking pictures every hour for 48 hours using infrared cameras and the software Pylon Viewer v5.1.0 (Basler©, Ahrensburg, Germany). For each tested male, we thus obtained an aggregation score ranging from 0 to 48, which was defined as the total number of pictures in which a male was in the chamber next to the group of conspecifics. After the 48h of aggregation test, we returned each male to its Petri dish until its use for subsequent measurements (see below). For ease of handling, all manipulated earwigs were anaesthetised with CO_2_ during the set-up of this last measurement.

### Sperm storage measurement

Of the 91 males still alive on day 31 after isolation (Figure 1), we used a random subset of 39 to measure sperm storage as the number of sperm present in their seminal vesicles (Figure 1). This counting occurred about two months after the males were separated from the females, which is probably long enough for the males to rebuild their sperm reserves, regardless of their previous mating rate. We counted sperm numbers following the protocol detailed in Damiens et al. (2002). In brief, we dissected each male under a dissecting microscope, placed their seminal vesicles on a slide with 15 µL of 1x Phosphate Buffered Saline and then pierced it to release all the sperm. We subsequently dried the plate on a heating block, after which the smear of sperm was sealed with a 70% ethanol solution and allowed to dry again at ambient temperature. The slides were then stored at 3°C. Two days later, we deposited 20µL of DAPI dye (concentration = 10 µmol/L) on the smear, covered it with a small piece of plastic wrap to allow the dye to infiltrate the cells while not drying out, and 10 minutes later, we replaced the plastic wrap with a glass slide cover. The slide was then viewed under the microscope at 20x magnification, and we took pictures of 5 different fields of the smear. Using an Olympus micrometre calibration slide and the software Image J (Schneider et al., 2012), we then counted the number of spermatozoa in all 5 fields for each male. Finally, we used all these numbers to calculate the number of spermatozoa per millimetre square and multiplied this number by the total area of the smear to obtain the total number of sperm per male. It should be noted that sperm storage was measured in males that survived 31 days in isolation without access to food, so that this value represents the sperm storage of individuals best adapted to these two stressful conditions.

### Survival in harsh environments and after exposure to pathogens

We measured male survival under two types of harsh conditions. The first type of harsh condition was the absence of any food source (starvation) combined with social isolation, which is known to have detrimental effects on this gregarious species (Kohlmeier et al., 2016; Van Meyel & Meunier, 2022). We assessed the male survival rate under these conditions by recording whether each of the 120 males tested was still alive on day 31 after isolation (Figure 1).

The second type of harsh condition was exposure to pathogens. We measured survival rate after pathogen exposure in the 52 males that were still alive on day 31 and were not used to measure sperm storage (Figure 1). We exposed each male to spores of the entomopathogenic fungus *Metharizium brunneum* (formerly *M. anisopliae*). This fungus is a natural and lethal pathogen of *F. auricularia* (Günther & Herter, 1974; Arcila & Meunier, 2020; Coulm & Meunier, 2021). The infection followed a standard protocol detailed in Kohlmeier et al. (2016). In brief, we immersed each male in an Eppendorf tube containing 1.5 mL of a conidiospore suspension of *M. brunneum* diluted in 0.05% Tween 20 (Sigma P-1379) at a concentration of 10^6^ spores/mL. We then gently swirled the tube from side to side for 4 seconds, removed the male and placed it back in its original Petri dish with standard food (see above) that was changed twice a week. We then kept the infected males at 20°C on a 12:12 light:dark schedule. We checked them daily for mortality over the course of 45 days. As with sperm storage, survival after pathogen exposure was measured in males that survived 31 days in isolation without access to food, so that it represents the survival rate of individuals best adapted to these two stressful conditions.

### Statistical analyses

We conducted all statistical analyses using the software R v4.3.2 (https://www.r-project.org/) loaded with the packages *DHARMa* (Hartig, 2020), *car* (Fox & Weisberg, 2019), *survival* (Therneau, 2020) and *emmeans* (Lenth, 2022). We analysed locomotor performance, sperm count and aggregation score using three general linear models (*lm* in R), while we analysed the likelihood to flee (yes or no) and whether males were dead 31 days after isolation (yes or no) using two generalized linear model (*glm* in R) with binomial error distributions. Finally, we analysed the survival rate of the pathogen-exposed males using a Cox proportional hazard regression model allowing for censored data (*Coxph* in R), i.e., males still alive 45 days after exposure to the pathogen. In these six models, we entered the type of forceps length (long or short), the population of origin of the males (Cinais or Valence) and the interaction between these two variables as explanatory factors. Overall, we checked that all model assumptions were met (e.g., homoscedasticity and normality of residuals) using the *DHARMa* package. To this end, we log-transformed male locomotor performance and log+1-transformed the aggregation scores. In the model where we found a significant interaction, we conducted pairwise comparisons using the estimated marginal means of the models and we corrected P values for multiple testing using the Tukey method, as implemented in the *emmeans* R package. Finally, we conducted power analyses for each of our statistical models using the packages *pscl* (Jackman et al., 2024), *pwr* (Champely et al., 2020) and *powerSurvEpi* (Qiu et al., 2021).

## Results

Overall, we detected no main association between forceps length and the six traits measured (Table 1). This applied to male locomotor performance, likelihood of fleeing after a physical disturbance, aggregation score, sperm storage and survival after pathogen infection (Figure 3; Table 1). There was also no difference between males with short and long forceps in terms of survival under harsh environmental conditions, although males with long forceps survived better when they came from Valence compared to Cinais (this trend was not present in males with short forceps; Figure 3; interaction in Table 1; pairwise comparisons: Long forceps Cinais vs Valence: Z = −3.311; P = 0.005; Short forceps Cinais vs Valence: Z = −0.611; P = 0.929). Regardless of forceps length, males from Valence were overall more likely to flee after a physical disturbance and less gregarious than males from Cinais (Table 1). These differences between the two populations were absent for all other traits measured (Table 1). There was no interaction between the population of origin and forceps length for any of the traits measured, except for survival under harsh environmental conditions (Table 1). The statistical power of each analysis ranges from 0.155 (sperm count) to 0.447 (survival in harsh environments; Table 1). These values suggest that the likelihood of detecting statistically significant effects based on the values reported in this study was sometimes relatively low, particularly given the large variance obtained for certain traits. Larger sample sizes, particularly for sperm count, may therefore be needed to confirm the absence of effects more robustly.

**Table 1.**
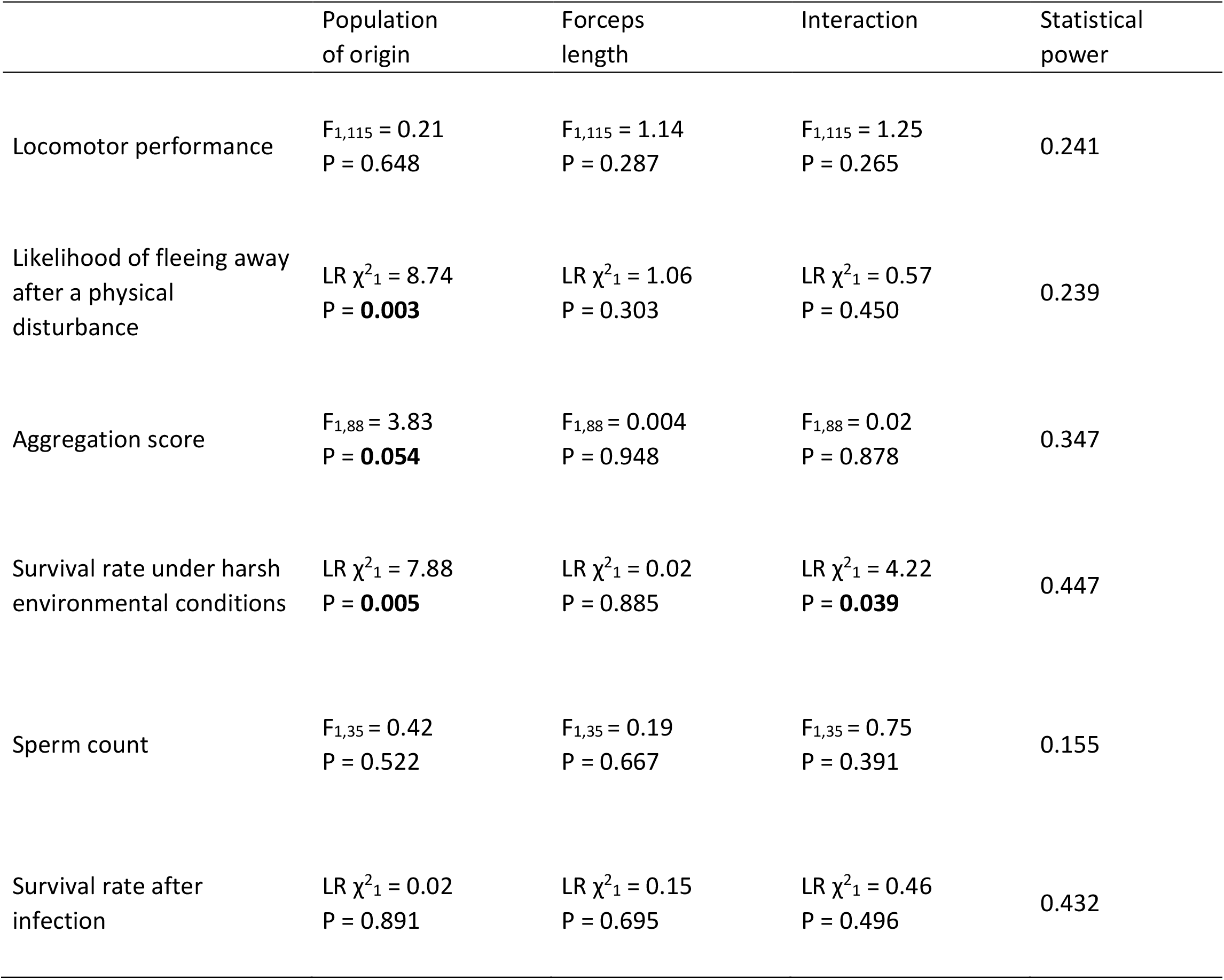
Effects of male forceps length and population of origin of the three physiological and three behaviours measured in this study. Significant p-values are in bold.

**Figure 3.**
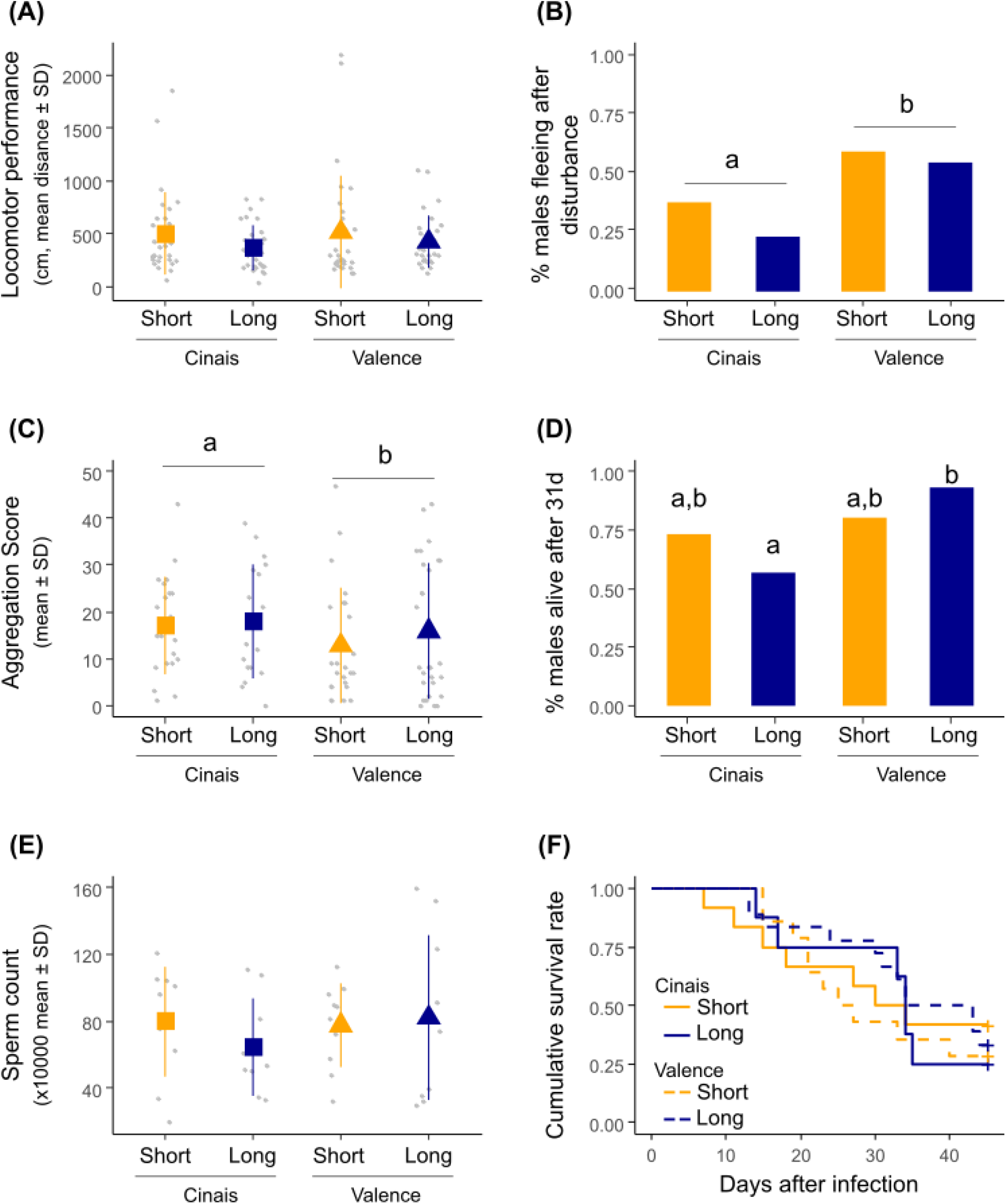
Association between forceps length and male (A) locomotor performance, (B) likelihood of fleeing after a physical disturbance, (C) propensity to aggregate with conspecifics, (D) likelihood of being alive after 31 days in social isolation and without food access, (E) sperm storage in the vesicle and (F) survival rate after pathogen infection. Different letters *P* <0.05.

## Discussion

The display of large, extravagant weapons often comes at a cost to the males (Emlen, 2008; McCullough et al., 2016; Goldberg et al., 2019). It is therefore expected that only males in the best condition can afford them and thus that these large weapons signal good male quality (Otte & Stayman, 1979; Emlen & Nijhout, 2000). Here, we tested this prediction in the European earwig by investigating whether male forceps length is associated with six important behavioural and physiological traits. Contrary to predictions, our experiment did not allow us to detect an association between forceps length and locomotor performance, boldness (i.e., likelihood to flee after a physical disturbance), aggregation behaviour, sperm production, and male survival after pathogen infection. These findings were consistent between the two populations, although some of the traits measured were population specific: males from Cinais were generally bolder, less gregarious and (only if they carried long-forceps) had a better chance of surviving in harsh conditions than males from Valence.

Our results contrast with much of the literature reporting associations between sexually selected male attributes and physiological, behavioural or immunological traits in arthropods (Emlen, 2008; McCullough et al., 2016; Goldberg et al., 2019). However, the European earwig is not the only case where such associations appear to be lacking (Emlen, 2008; Swallow & Husak, 2011). For example, carrying giant claw is not associated with the efficiency of escape behavior and the level of metabolic costs in two fiddler crabs (Tullis & Straube, 2017; Pena & Levinton, 2021). Similarly, bearing large horns does not reflect male growth, mobility, or immunity in the rhinoceros beetle *Trypoxylus dichotomus* (McCullough & Emlen, 2013; McCullough & Tobalske, 2013). It has been suggested that this apparent lack of general association may be due to the fact that the performance of each weapon size depend on the environment in which the weapons are used and/or because weapon sizes reflect alternative reproductive tactics (McCullough & Emlen, 2013; McCullough et al., 2016). This could be the case with the European earwig (Tomkins et al., 2005). In this species, males with large forceps have been suggested to be more active in guarding of females, while males with small forceps are more active in sneaking into copulations (Tomkins & Brown, 2004; Kamimura, 2014). Having long forceps should therefore be rewarded more frequently when the encounter rate between competitors is high (Hunt & Simmons, 2001), such as in high population densities. Consistent with this prediction, Tomkins & Brown (2004) found that the proportion of long forceps males increased with population densities across 46 island and mainland sites in the UK. Another possible explanation for our results is that the association between male forceps length and the six traits measured could have been masked by differences in the amount of resources available to each male and/or used by each male for these traits (van Noordwijk & de Jong, 1986). In our study, all males were kept under identical laboratory conditions and fed *ad libitum* for three months prior to our measurement. This explanation would therefore suggest that the variation in resources influencing investment decisions is determined prior to field sampling, e.g. during development or early adulthood. Previous data suggest that it could be the case in the European earwig. In this species, early life conditions have long-term effects on the physiology and behaviour of adults (Wong & Kölliker, 2014; Thesing et al., 2015; Raveh et al., 2016) and inter-individual variation in female condition can mask investment trade-off between egg quantity and quality (Koch & Meunier, 2014). Overall, our findings call for future studies to confirm whether males with short or long forceps have alternative reproductive tactics, whether their reproductive success depends on population densities and whether early life conditions can affect the likelihood to detect the association between forceps length and the six traits measured. They also call for further research to quantify other potential costs of carrying long forceps in this species, for example in terms of predation rates and ability to fly (Crumb & Eide, 1941). These notwithstanding, our findings emphasize that forceps size is not a good predictor of the six behavioural and physiological traits measured in males.

While we found no overall difference between short- and long-forceps males, our data reveal population differences in terms of males’ boldness, aggregation level and resistance to starvation. Cinais had males that were generally bolder, more gregarious, as well as males with long forceps that survived food deprivation less well than Valence. These effects are unlikely to reflect a plastic response to the direct environment of the males tested (e.g. population differences in terms of nutritional status), as they were all reared under common laboratory conditions for months prior to the start of our experiments. Instead, it could reflect population idiosyncrasies that have affected their development, such as climatic conditions (Valence is warmer than Cinais), environmental conditions and/or exposure to phytosanitary products (e.g. Valence is a cultivated orchard, whereas Cinais is an uncultivated forest edge), or population-specific genetic background. In line with this hypothesis, several life history traits of the European earwig are known to be shaped by the duration of cold during egg development, the level of warm temperatures during nymph development and, more generally, by seasonal parameters encountered by offspring during development (Körner et al., 2018; Tourneur & Meunier, 2020; Coulm & Meunier, 2021). Similarly, recent studies have shown that the European earwig can be sensitive to exposure to even low levels of pesticides, which can lead to populations specificities in terms of earwig physiology, morphology and behaviour (Malagnoux, Capowiez, et al., 2015; Malagnoux, Marliac, et al., 2015; Le Navenant et al., 2019; Meunier et al., 2020; Mauduit et al., 2021; Fricaux et al., 2023). Regardless of the nature of these population idiosyncrasies, our study shows that they do not affect the (lack of) association between forceps length and the six traits measured.

Overall, our findings questions the robustness of our understanding of forceps length diversity in terms of function, maintenance and use as a possible quality signal in the European earwig (McCullough et al., 2016). To date, our knowledge of these questions is exclusively based on laboratory experiments in which females were presented to one or two males (reviewed in Kamimura, 2014). While the results of these studies suggest that males can gain fitness benefit from growing longer forceps (Radesäter & Halldórsdóttir, 1993; Styrsky & Rhein, 1999; Forslund, 2000, 2003; Walker & Fell, 2001), they contain two important limitations. First, they considered mating success (i.e., success in gaining copulation) and not reproductive success (i.e. success in producing offspring). As these two parameters are not necessarily related (Thompson et al., 2011), it cannot be excluded that short forceps males have a similar or even higher reproductive output than their counterparts. This could be because mating is more efficient in short compared to long forceps males (Brown, 2006), or because short forceps males have alternative reproductive tactics to long forceps males (see above; Tomkins & Brown, 2004). One study used a genetic approach to study reproductive success and found no association between male forceps length and the number of offspring sired (Sandrin et al., 2015). However, this study only focused on length variation within short forceps males. The second limitation is that all these studies have examined mating success in groups of a maximum of two males. This number is far less than the hundreds of individuals that typically constitute earwig aggregates (Meunier, 2024a) and thus cannot rule out the possibility that the apparent advantage of long-forceps males in pairs or trios may disappear or even become a disadvantage under different demographic conditions (Hunt & Simmons, 2001; Tomkins & Brown, 2004; Oliveira et al., 2008). Hence, our current understanding of the relationship between forceps length and male fitness should be treated with caution in the European earwig. Our results thus call for future studies to determine the function and reliability of male forceps length under natural conditions, and the evolutionary drivers that explain the maintenance of its diversity within and between populations.

## Acknowledgments

We thank Lucas Marie-Orléach for his comments on a previous version of this manuscript. We also thank Jean-Christophe Lenoir and Armand Guillermin for their help with field sampling, and the INRAE unité expérimentale Recherche Intégrée Gotheron for giving us access to their orchards for earwig field sampling. Finally, we thank the two anonymous reviewers, Luna Grey and Olivier Roux for their comments on this manuscript.

Preprint version 3 of this article has been peer-reviewed and recommended by Peer Community In Zoology https://doi.org/10.24072/pci.zool.100318 (Roux, 2024)

## Funding

This project has been funded by the French National Research Agency (ANR-20-CE02-0002 to J.M).

## Conflict of interest disclosure

The authors declare that they have no financial conflict of interest with the content of this article. J.M is a recommender for PCI Zoology.

## Data, script and code availability

Data and R script are available online: https://doi.org/10.5281/zenodo.11469628 (Meunier, 2024b)

## Notes

https://doi.org/10.5281/zenodo.11469628

### Competing Interest Statement

The authors have declared no competing interest.

### Summary of Updates

Manuscript formatted according to Peer Community In Zoology requirements.

